# TRIP6 condensates constrain Hippo pathway hyperactivation under osmotic stress

**DOI:** 10.64898/2026.10.16.759540

**Authors:** Weihong Fu, Yuxi Yang, Lin Xie, Junying Li, Hua Li, Wenxiu Ning

## Abstract

The Hippo pathway integrates mechanical and chemical cues to regulate cell proliferation and tissue homeostasis, whereas its dysregulation contributes to impaired regeneration and tumorigenesis. Phase separation has emerged as an organizing principle for Hippo signal transduction, but how cells constrain excessive pathway activation under stress remain poorly defined. Here we demonstrate that TRIP6, a LIM-domain scaffold and tension-sensitive Hippo inhibitor, undergoes intrinsic phase separation both *in vitro* and in living cells. Osmotic stress triggers rapid redistribution of TRIP6 condensates from large cortical assemblies to dispersed cytoplasmic puncta. Domain mapping shows that the N-terminal IDR2 drives TRIP6 phase separation, whereas the C-terminal LIM domains negatively regulate condensate fluidity. Mechanistically, TRIP6 condensates selectively recruit the LATS1 inhibitor AJUBA and the MST1 scaffold SAV1 while excluding MST1 itself. The phase separation-deficient ΔIDR2 mutant fails to undergo stress-induced condensate redistribution, exhibits weakened LATS1 binding, and shows increased YAP phosphorylation and cytoplasmic retention under osmotic stress, with reduced cell proliferation and stress resilience. Finally, TRIP6 condensates disassemble in a mouse model of elevated epidermal tissue tension, identifying them as mechanosensitive platforms that link phase transitions to Hippo-dependent growth control. Together, these findings establish TRIP6 condensates as a negative regulatory node that buffers excessive Hippo activation to preserve cellular homeostasis under hyperosmotic conditions.

## Introduction

The Hippo signaling pathway is a conserved regulatory network that coordinates cell proliferation, tissue homeostasis and organ regeneration by integrating mechanical, chemical and metabolic cues [1]. Its dysregulation contributes to tumorigenesis, organ fibrosis and defective tissue repair, underscoring its broad biological and therapeutic significance [2]. Biomolecular condensation represents a core physicochemical mechanism for assembling membraneless condensates. These compartments locally enrich signaling components and tune the rates of intracellular biochemical reactions [3, 4]. Condensates possess adjustable material properties. They can undergo reversible assembly-disassembly upon diverse cellular stimuli, such as osmotic perturbations, mechanical force and post-translational modifications [5, 6]. Accumulating evidence establishes phase-separated assemblies as general modulators of major signal transduction pathways, providing spatiotemporal control over signal initiation, amplification and termination [7, 8]. This principle is particularly evident in the Hippo pathway, where phase separation governs signal transduction at multiple levels.

Multiple Hippo pathway components undergo phase separation at discrete signaling nodes. Upstream scaffold proteins nucleate MST-Hpo kinase condensates to promote kinase autophosphorylation and pathway activation [9], while the downstream effector YAP forms nuclear condensates whose disassembly by mechanical stress rewires transcriptional programs [10]. In stratified epithelia, RIPK4 drives non-canonical Hippo activation by recruiting LATS1/2 kinases into ankyrin repeat-mediated condensates [11]. Osmotic stress is a conserved physiological trigger for Hippo pathway activation, and phase separation-based mechanisms underpin this response. Osmotic challenge induces multiphase coalescence of pro-activating AMOT/KIBRA condensates with inhibitory SLMAP-STRIPAK condensates, which inactivates the phosphatase complex to fine-tune signaling output [12]. The tumor suppressor Merlin also forms PI4P-dependent solid-like apical condensates that serve as stable signaling platforms and confer resistance to hypoosmotic stress [13]. Despite these advances, molecular mechanisms that constrain excessive Hippo pathway activation to preserve cellular homeostasis under hyperosmotic conditions remain poorly defined.

LIM-domain-containing proteins represent a large superfamily unified by conserved tandem zinc-finger LIM modules [14, 15]. Many LIM family members reside at cell-cell adherens junctions and focal adhesions, acting as mechanosensitive molecular scaffolds. They convert mechanical stimuli into cytoskeletal rearrangement and downstream transcriptional signaling cascades [16, 17]. Evolutionarily conserved LIM-domain-dependent mechanisms enable detection of force-remodelled cytoskeletal filaments. This capability broadens the range of cellular mechanosensory strategies [18, 19]. These proteins assemble extensive multiprotein interactomes to execute tissue-specific physiological functions under mechanical stress conditions [20]. TRIP6 is a LIM-domain scaffold protein enriched at adherens junctions. It functions as a tension-sensitive negative regulator of Hippo signaling by directly binding LATS1/2 and outcompeting MOB1, thereby blocking MST1/2 recruitment [21]. Subsequent work has further placed the TRIP6-LATS1 complex as an integral component of the α-catenin/vinculin tension sensor apparatus at both bicellular and tricellular junctions where its junctional localization is strictly dependent on actomyosin contractility [22]. Notably, LIMD1, another LIM-domain protein, forms force-sensitive condensates at focal adhesions, and TRIP6 has been identified as a client protein enriched within these condensates [23]. This prompted us to ask whether TRIP6 itself possesses intrinsic phase separation capacity and how its condensates modulate Hippo pathway under osmotic stress.

In this study, we demonstrate that TRIP6 undergoes phase separation both *in vitro* and in living cells, and that osmotic stress triggers rapid redistribution of TRIP6 condensates from large cortical assemblies to dispersed cytoplasmic puncta. We further show that IDR2-mediated phase separation enables TRIP6 to act as an attenuator of excessive Hippo pathway activation under hyperosmotic conditions, thereby preserving cell proliferation and stress resilience.

## Results

### TRIP6 restrains Hippo hyperactivation and intrinsically forms condensates

Sorbitol-induced hyperosmotic stress is a well-validated trigger for Hippo activation and promotes condensate formation of upstream Hippo regulators [12]. The subcellular response of TRIP6 to hyperosmotic challenge remains uncharacterized. We thus first profiled the subcellular dynamics of endogenous TRIP6 in DLD1 cells in response to osmotic stress. Under basal culture conditions, TRIP6 predominantly accumulated at the cell cortex, forming large discrete punctate assemblies aligned with cortical F-actin bundles and stress fibers. Upon treatment with 0.4 M sorbitol, these large cortical structures disassembled, and TRIP6 reorganized into numerous small, uniformly distributed condensates scattered throughout the cytoplasm. This redistribution was fully recapitulated by 0.2 M NaCl, an independent hyperosmotic stimulus, indicating that TRIP6 redistribution is a general cellular response to hyperosmotic challenge (Figure 1A). We then knocked out endogenous TRIP6 in DLD1 cells via CRISPR-Cas9, and confirmed the knockout efficiency by western blot and immunofluorescence staining (Figure 1B-C). We next evaluated the effect of TRIP6 on Hippo pathway activity under osmotic stress, with YAP phosphorylation at Ser127 as the canonical readout. Compared with scramble control cells, TRIP6 KO cells displayed markedly elevated YAP phosphorylation with a clear electrophoretic mobility shift at 30-60 min after sorbitol treatment, indicating that TRIP6 restrains excessive Hippo pathway activation under osmotic stress (Figure 1D). Consistent with this regulatory role, loss of TRIP6 impaired cellular stress resilience. CCK-8 assays showed that TRIP6 KO cells exhibited a more pronounced decrease in viability than controls following prolonged sorbitol treatment (Figure 1E). TRIP6 deletion also suppressed clonogenic growth, with a marked reduction in colony area (Figure 1F-G). To better characterize the features of TRIP6 condensates, we generated a stable TRIP6-GFP-expressing cell line in the TRIP6-knockout DLD1 background. Western blot confirmed the expression of TRIP6-GFP (Figure S1A). Immunofluorescence staining showed that TRIP6-GFP similarly underwent subcellular redistribution in response to osmotic stress (Figure S1B). Structured illumination microscopy (SIM) imaging revealed that under basal culture conditions, TRIP6-GFP formed large condensates distributed along cortical F-actin fibers. Upon stimulation with 0.4 M sorbitol, these large cortical structures disassembled and reorganized into smaller, uniformly distributed condensates throughout the cytoplasm (Figure 1H). We performed fluorescence recovery after photobleaching (FRAP) assays on these large TRIP6-GFP condensates. The fluorescence intensity recovered to approximately 40% of the pre-bleaching level, indicating that the condensates possess moderate molecular mobility (Figure 1I-J).

**Figure 1.**
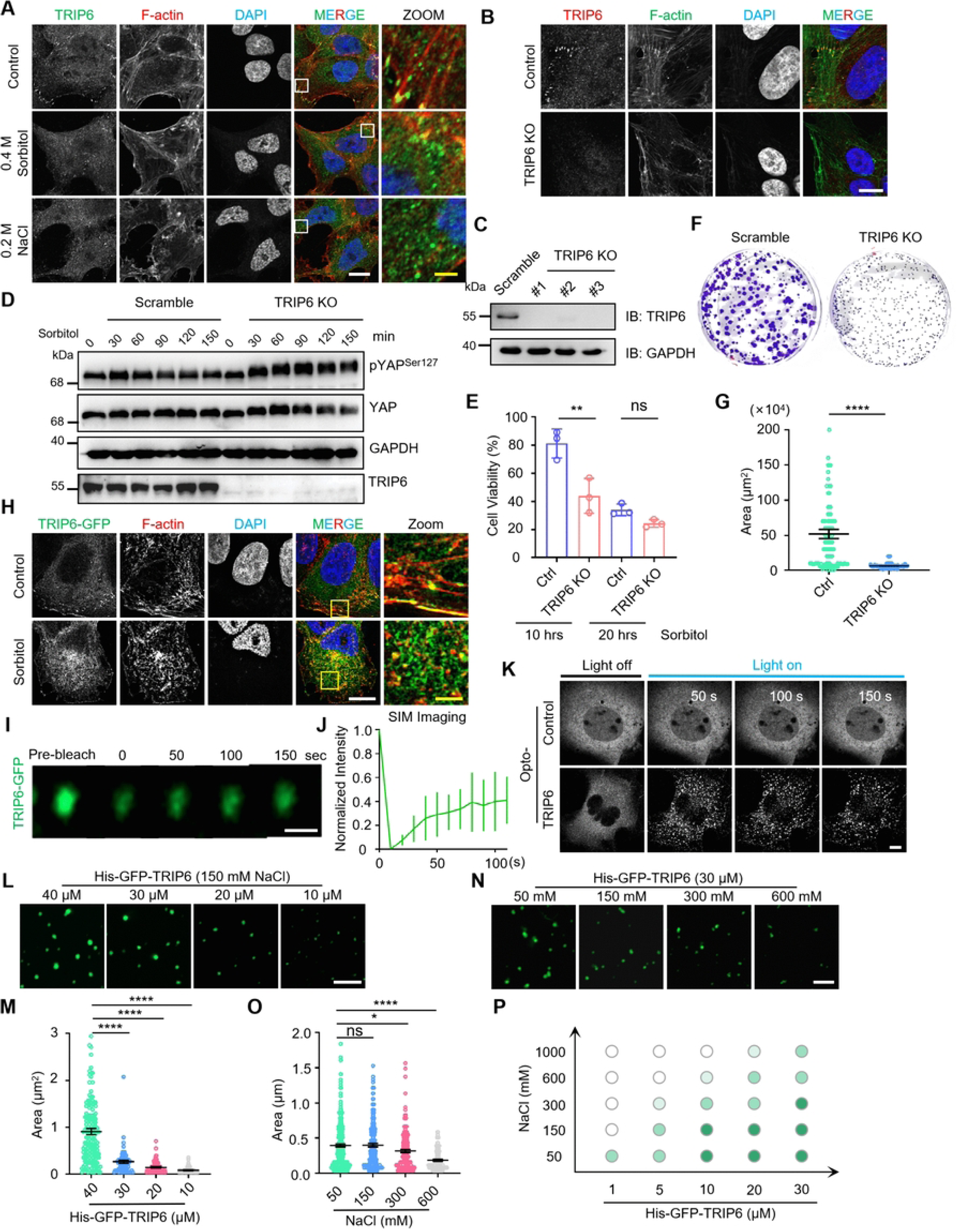
TRIP6 restrains Hippo hyperactivation and intrinsically forms condensates. (A) Immunofluorescence staining of endogenous TRIP6 in DLD1 cells under basal conditions, 0.4 M sorbitol and 0.2 M NaCl treatment. TRIP6 (green), F-actin (red), DAPI (blue). TRIP6 redistributes from cortical puncta to dispersed cytoplasmic condensates upon hyperosmotic challenge. Scale bar, 10 μm (white) and 2 μm (yellow). (B and C) Validation of TRIP6 knockout in DLD1 cells. (B) Immunofluorescence staining confirming the absence of TRIP6 signal in TRIP6 KO cells. Scale bar, 10 μm. (C) Western blot analysis of TRIP6 protein levels in scramble and three independent TRIP6 KO clones. (D) Western blot analysis of YAP phosphorylation at Ser127 in scramble control and TRIP6 knockout (KO) DLD1 cells treated with 0.4 M sorbitol for the indicated times. TRIP6 KO cells showed elevated YAP phosphorylation with a mobility shift at 30-60 min. (E) CCK-8 assay showing that TRIP6 KO cells exhibit a more pronounced decrease in viability than controls after prolonged sorbitol treatment. n = 3 independent experiments per group. Data are presented as mean ± SD. One-way ANOVA followed by Tukey’s multiple comparisons test. ns, not significant, **p < 0.01. (F and G) Clonogenic assays showing reduced colony formation in TRIP6 KO cells. (F) Representative images of crystal violet-stained colonies. (G) Quantification of colony area. n = 100 colonies per group. Data are presented as mean ± SEM. Unpaired t test. ****p < 0.0001. (H) Super-resolution microscopy (SIM) images of TRIP6-GFP cells under basal conditions or after 0.4 M sorbitol treatment. TRIP6-GFP (green) formed large condensates along cortical F-actin fibers (red) under basal conditions and redistributed into smaller dispersed cytoplasmic condensates upon sorbitol stimulation. DAPI (blue) marks nuclei. Scale bars, 10 μm (white) and 2 μm (yellow). (I and J) FRAP analysis of TRIP6-GFP condensates. (I) Representative time-lapse images before and after photobleaching. Scale bar, 1 μm. (J) Quantification of fluorescence recovery showing approximately 40% recovery. n = 4 independent experiments. Data are mean ± SD. (K) OptoDroplet assay showing light-induced condensate formation by Opto-TRIP6, but not the control construct. Scale bar, 10 μm. (L and M) *In vitro* phase separation of purified His-GFP-TRIP6 at the indicated concentrations in 150 mM NaCl. (L) Representative images. Scale bar, 5 μm. (M) Quantification of droplet area showing concentration-dependent phase separation. n = 168 (40 μM), 68 (30 μM), 86 (20 μM), 63 (10 μM) droplets, respectively. Data are presented as mean ± SEM. One-way ANOVA followed by Dunnett’s multiple comparisons test. ****p < 0.0001. (N) Representative images of TRIP6 droplets at the indicated NaCl concentrations. Scale bar, 5 μm. (O) Quantification of droplet area showing progressive reduction with increasing NaCl concentration. n = 284 (50 mM), 68 (150 mM), 150 (300 mM), 98 (600 mM) droplets, respectively. Data are presented as mean ± SEM. One-way ANOVA followed by Dunnett’s multiple comparisons test. ns, not significant; *p < 0.05, ****p < 0.0001. (P) Phase diagram of TRIP6 across a range of protein and NaCl concentrations.

Since osmotic-stress induced redistribution of TRIP6 condensates, we wondered whether TRIP6 possesses an intrinsic capacity for phase separation independent of cellular cofactors. We employed the optoDroplet system, a blue-light inducible platform for spatiotemporal control of intracellular phase separation in living cells [24]. TRIP6 exhibited diffuse cytoplasmic distribution without blue-light activation, while it rapidly assembled into numerous discrete cytoplasmic puncta within 50 seconds after blue-light illumination. No puncta formation was observed in control cells under identical illumination conditions (Figure 1K). These data demonstrate that induced TRIP6 oligomerization is sufficient to drive phase separation in the intracellular environment. To establish whether TRIP6 phase separation is an intrinsic biophysical property, we expressed and purified recombinant His-GFP-tagged TRIP6 from *Escherichia coli* (Figure S1C). Under physiological ionic strength (150 mM NaCl), purified TRIP6 spontaneously formed spherical, micron-scale droplets and exhibited concentration-dependent behavior, with condensate size increasing progressively as protein concentration rose from 10 μM to 40 μM (Figure 1L-M). As biomolecular phase separation is generally driven by weak multivalent intermolecular interactions and highly sensitive to ionic strength, we further assessed the effect of NaCl concentration on TRIP6 phase behavior at a fixed protein concentration of 30 μM. Increasing NaCl from 50 mM to 600 mM led to a progressive reduction in both the size and number of TRIP6 droplets (Figure 1N-O). A systematic phase diagram spanning a range of protein and salt concentrations confirmed that TRIP6 phase separation is cooperatively modulated by both parameters, with higher protein concentration and lower ionic strength favoring formation of the condensed phase (Figure 1P). Collectively, these data demonstrate that TRIP6 is an intrinsically phase-separating protein whose condensation behavior is tunable by both protein concentration and ionic environment, providing a biophysical basis for its osmotic stress-dependent redistribution in cells.

### LIM domains tune condensate fluidity and IDR2 drives TRIP6 condensation

We next dissected the structural basis of TRIP6 phase separation. Sequence-based disorder prediction revealed two intrinsically disordered regions (named IDR1 and IDR2) in the N-terminal half of TRIP6, whereas the C-terminal ordered regions harbor three tandem LIM domains and a PDZ domain (Figure 2A). We sought to determine whether its phase separation capacity is encoded by the disordered N-terminal region or the structured C-terminal region. *In vitro* phase separation assays using purified recombinant proteins showed that both TRIP6 wild type (WT) and the N-terminal fragment spontaneously formed spherical droplets, with the N-terminal yielding significantly larger droplets, while the C-terminal exhibited no phase separation activity at the same concentration (Figures S2A, 2B-C). The optoDroplet system further confirmed these findings in living cells, showing that blue-light illumination induced robust condensate formation by TRIP6 WT and the N-terminal, whereas the C-terminal remained diffusely distributed (Figure 2D). To further determine whether the C-terminal LIM domains regulate TRIP6 phase separation, we generated and purified sequential LIM deletion mutants (ΔLIM3, ΔLIM2/3, ΔLIM1/2/3) (Figure S2B). *In vitro* phase separation assays showed that all mutants underwent phase separation, with no significant difference in droplet area compared with TRIP6 WT (Figure S2C-D). We then performed FRAP assays to compare condensate fluidity across these mutants. TRIP6 WT condensates showed minimal fluorescence recovery, indicating very low intrinsic fluidity. Interestingly, the recovery rate and mobile fraction of the LIM domain deleted mutations increased progressively, with ΔLIM1/2/3 condensates showing the fastest and most complete recovery (Figure 2E-F). Treatment with 1,6-hexanediol, which disrupts weak hydrophobic intermolecular interactions, reduced condensate area across all LIM deleted mutants, and the extent of dissolution increased with the number of LIM domains removed (Figure 2G-H). Time-lapse imaging showed that adjacent TRIP6 WT condensates rarely underwent fusion, whereas ΔLIM1/2/3 droplets fused readily over time (Figure 2I). Collectively, these results demonstrate that LIM domains exert a stepwise negative regulatory effect on the fluidity of TRIP6 condensates, stabilizing them in a less dynamic state.

**Figure 2.**
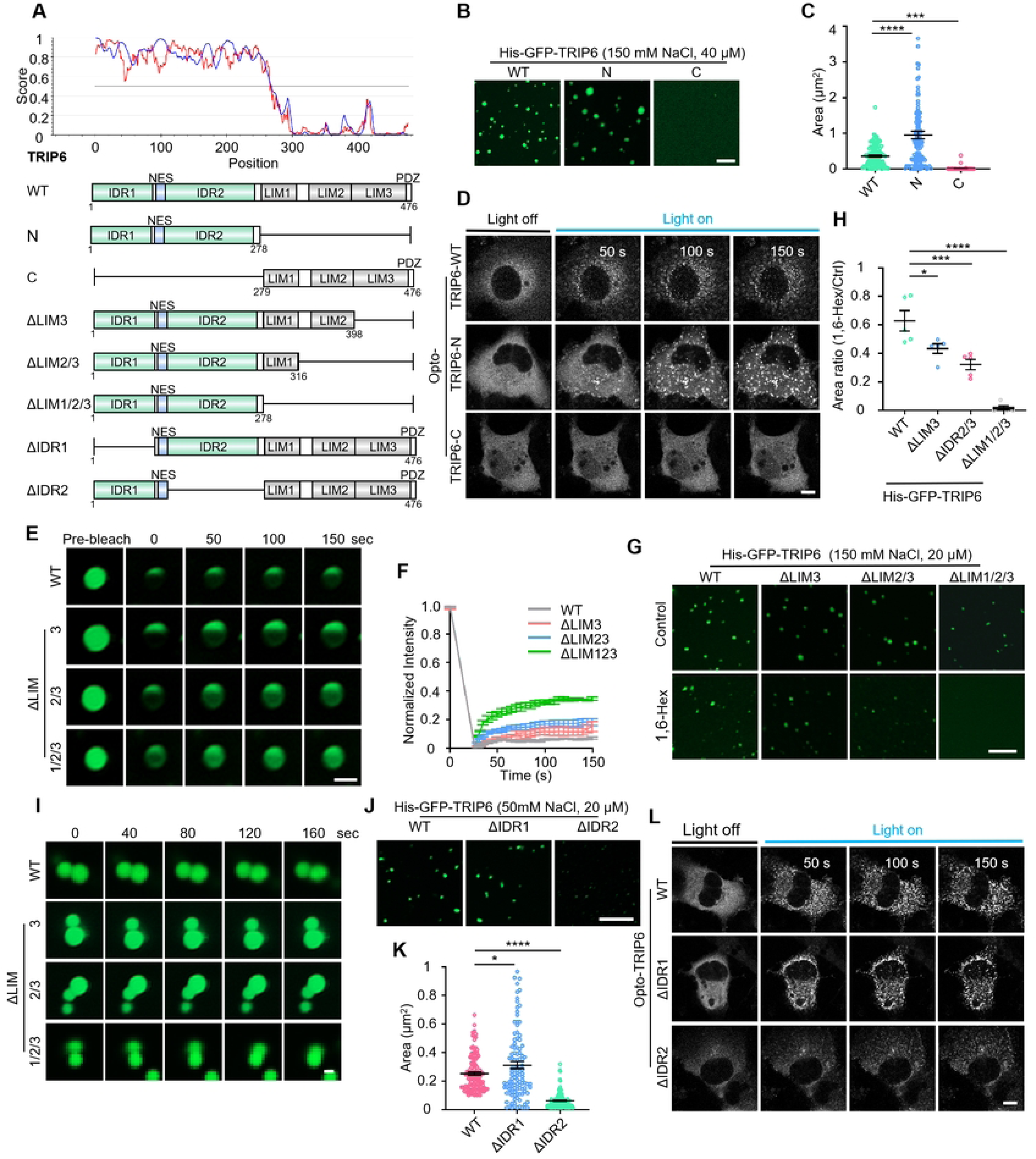
LIM domains tune condensate fluidity and IDR2 drives TRIP6 condensation. (A) Sequence-based disorder prediction and domain architecture of TRIP6 used in this study, showing two N-terminal IDRs (IDR1 and IDR2) and C-terminal LIM and PDZ domains. (B and C) *In vitro* phase separation of TRIP6 (WT), N-terminal (N), and C-terminal (C) fragments at 40 μM in 150 mM NaCl. (B) Representative images. Scale bar, 5 μm. (C) Quantification of droplet area. n = 120 droplets per group, respectively. Data are presented as mean ± SEM. One-way ANOVA followed by Tukey’s multiple comparisons test. ****p < 0.0001, ***p < 0.001. (D) OptoDroplet assay of WT, N-terminal and C-terminal TRIP6 constructs showing that blue light induces condensate formation by WT and the N-terminal fragment but not the C-terminal fragment. Scale bar, 10 μm. (E and F) FRAP analysis of WT and sequential LIM deletion mutants. (E) Time-lapse images of WT, ΔLIM3, ΔLIM2/3 and ΔLIM1/2/3 condensates. Scale bar, 1 μm. (F) Quantification showing progressive increase in fluorescence recovery with successive LIM domain deletions. n = 3 independent experiments per group, respectively. Data are presented as mean ± SEM. (G and H) 1,6-hexanediol sensitivity of WT and LIM deletion mutant condensates. (G) Representative images before and after 1,6-hexanediol treatment. Scale bar, 5 μm. (H) Quantification of condensate area ratio after 1,6-hexanediol treatment showing increased dissolution with successive LIM domain deletions. n = 5 independent experiments for each group. Data are presented as mean ± SEM. One-way ANOVA followed by Tukey’s multiple comparisons test. *p < 0.05, ***p < 0.001, ****p < 0.0001. (I) Time-lapse imaging of droplet fusion for WT, ΔLIM3, ΔLIM2/3 and ΔLIM1/2/3 condensates over 160 s. WT condensates rarely fuse, whereas ΔLIM1/2/3 droplets coalesce efficiently over time. Scale bar, 1 μm. (J and K) *In vitro* phase separation of WT, ΔIDR1 and ΔIDR2 mutants at 20 μM in 50 mM NaCl. (J) Representative images. Scale bar, 5 μm. (K) Quantification of droplet area showing that ΔIDR2 forms only a few very small condensates. n = 117 (WT), 113 (ΔIDR1), 117 (ΔIDR2) droplets, respectively. Data are presented as mean ± SEM. One-way ANOVA followed by Tukey’s multiple comparisons test. *p < 0.05, ****p < 0.0001. (L) OptoDroplet assay of WT, ΔIDR1 and ΔIDR2 mutants showing that ΔIDR2 is severely impaired in light-induced condensate formation. Scale bar, 10 μm.

Building on the finding that the N-terminal region drives TRIP6 phase separation, we sought to determine which of its two disordered regions, IDR1 or IDR2, is the key determinant. We thus generated and purified IDR1 and IDR2 deletion mutants (ΔIDR1 and ΔIDR2). *In vitro* phase separation assays showed that ΔIDR1 retained phase separation activity comparable to TRIP6 WT, whereas ΔIDR2 formed only a few very small condensates (Figures S3A, 2J-K). Consistently, in the optoDroplet assay, blue-light illumination induced robust condensate formation by TRIP6 WT and ΔIDR1, whereas ΔIDR2 formed only sparse, small puncta even after prolonged light stimulation (Figure 2L). These results identify IDR2 as the major driver of TRIP6 phase separation. To determine whether phase separation activity of the IDR2 arises from its intrinsic disorder rather than its specific amino acid sequence, we replaced IDR2 with the well-characterized IDR from FUS (Figure S3B) [25]. The optoDroplet assay showed that this FUS-IDR substitution restored light-inducible condensate formation in living cells (Figure S3C). FUS-IDR also formed large droplets at a concentration as low as 10 μM *in vitro*, indicating enhanced phase separation capacity (Figure S3D-F). This rescue by a heterologous IDR demonstrates that the intrinsic disorder of IDR2 serves as the core driving force for TRIP6 phase separation.

### TRIP6 condensates selectively partition Hippo pathway components and mediate osmotic stress-enhanced association with LATS1

Having established the biophysical basis of TRIP6 phase separation, we next sought to define the molecular constituents of TRIP6 condensates in cells using TurboID-mediated proximity labeling [26], which enables covalent biotinylation of proximal proteins within condensates in living cells (Figure 3A). We constructed C-terminally TurboID-Flag (FT)-tagged TRIP6 (TRIP6 WT-FT) and the phase separation-deficient TRIP6 ΔIDR2-FT mutant, with free TurboID FT as a negative control. Streptavidin pull-down assays followed by western blot analysis confirmed robust expression and efficient biotinylated protein enrichment for all three fusion constructs (Figure S4A). Immunofluorescence staining further confirmed efficient expression of these fusion constructs in DLD1 cells (Figure S4B). We then subjected the enriched proteins to quantitative mass spectrometry. Principal component analysis of the resulting quantitative proteomic dataset revealed clear separation among control, TRIP6 WT and TRIP6 ΔIDR2 samples with excellent intra-group reproducibility (Figure S4C). KEGG pathway enrichment analysis identified the Hippo signaling pathway as the most significantly enriched functional category in TRIP6 WT condensates, along with tight junction, adherens junction, endocytosis and actin cytoskeleton regulation (Figure 3B). Specifically, core components and regulators of the Hippo pathway including AJUBA, MST1 and SAV1, were substantially enriched in TRIP6 WT samples compared with the ΔIDR2 mutant and control groups (Figure 3C). To further validate whether these three proteins are recruited into TRIP6 condensates, we tagged AJUBA, SAV1 and MST1 with His-mCherry, purified the recombinant proteins, and performed *in vitro* co-phase separation assays with TRIP6-GFP. AJUBA and SAV1 efficiently co-condensed with TRIP6-GFP droplets, whereas MST1 did not partition into TRIP6 condensates (Figures 3D-G, S4D). Co-expression experiments in HeLa cells further confirmed these findings (Figure 3H-K). Together, these data indicate that TRIP6 condensates selectively recruit the LATS1 inhibitor AJUBA and the MST1 scaffold protein SAV1 while excluding MST1, suggesting that they may modulate Hippo signaling pathway by regulating LATS1 activation.

**Figure 3.**
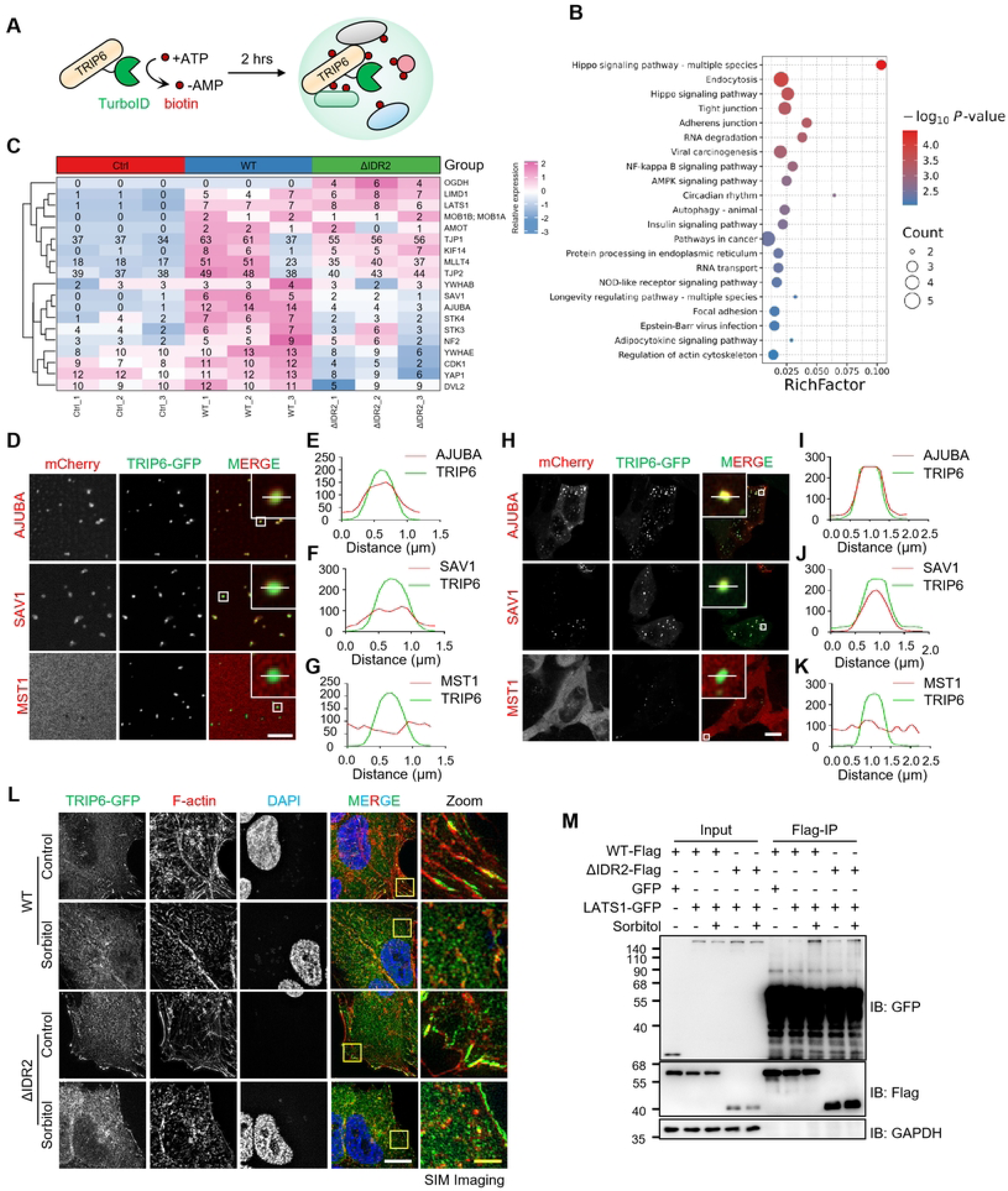
TRIP6 condensates selectively recruit Hippo regulators and mediate stress-enhanced TRIP6-LATS1 interaction. (A) Schematic illustration of TurboID-mediated proximity labeling to identify constituents of TRIP6 condensates in living cells. (C) Hierarchical clustering heatmap showing the enrichment of core Hippo pathway components in TRIP6 WT, ΔIDR2, and control samples as determined by quantitative mass spectrometry. (D-G) *In vitro* co-phase separation assays of TRIP6-GFP with mCherry-tagged AJUBA, SAV1 and MST1. (D) Representative images showing that AJUBA and SAV1 co-condense with TRIP6-GFP droplets, whereas MST1 remains excluded. Scale bar, 5 μm. (E-G) Fluorescence intensity line-scan profiles across individual droplets confirming co-localization of AJUBA and SAV1 with TRIP6 and the exclusion of MST1. F.I. fluorescence intensity. (H-K) Co-expression experiments in HeLa cells confirming co-localization of AJUBA and SAV1 with TRIP6-GFP condensates and exclusion of MST1. (H) Representative images. Scale bar, 10 μm. (I-K) Fluorescence intensity line-scan profiles across individual condensates. F.I., fluorescence intensity. (L) SIM images of TRIP6-GFP WT and ΔIDR2 cells under basal conditions and after sorbitol treatment. TRIP6 WT efficiently disassembles cortical condensates and redistributes to the cytoplasm upon sorbitol stimulation, whereas ΔIDR2 retains a substantial cortical pool. Scale bars, 10 μm (white) and 2 μm (yellow). (M) Co-immunoprecipitation assays showing that sorbitol treatment selectively enhances the interaction between TRIP6 WT and LATS1, an effect not observed in TRIP6 ΔIDR2 cells.

Since TRIP6 condensates undergo localization redistribution under osmotic stress and restrain excessive Hippo pathway activation, we next asked whether this regulatory function depends on TRIP6 phase separation. To address this, we generated stable cell lines re-expressing TRIP6 WT or the phase separation-deficient ΔIDR2 mutant in the TRIP6-knockout background, and enriched positive cells by flow cytometry (Figure S5A-B). SIM imaging showed that upon sorbitol treatment, cortical TRIP6 condensates efficiently disassembled and redistributed into smaller cytoplasmic assemblies. In contrast, TRIP6 ΔIDR2 retained a substantial cortical pool, indicating that its condensate redistribution is impaired (Figure 3L). These results establish that IDR2-mediated phase separation is required for efficient stress-induced structural reorganization of TRIP6 condensates. However, how this impaired condensate redistribution affects Hippo pathway regulation remains unclear. Given that TRIP6 regulates the Hippo pathway by binding and inhibiting LATS1 [21], we asked whether the TRIP6-LATS1 interaction is regulated by TRIP6 phase separation. Co-immunoprecipitation assays showed that both TRIP6 and TRIP6 ΔIDR2 associated with LATS1 under basal conditions. Upon sorbitol treatment, the interaction between TRIP6 WT and LATS1 was markedly enhanced, whereas the ΔIDR2 mutant showed only a marginal increase (Figure 3M). Immunofluorescence staining further supported these findings, showing that sorbitol treatment markedly enhanced the colocalization between TRIP6 WT and LATS1, but not between TRIP6 ΔIDR2 and LATS1 (Figure S5C-G). These findings indicate that TRIP6 condensates act as dynamic scaffolding platforms that strengthen interactions with LATS1 in response to osmotic stress in an IDR2-dependent manner.

### IDR2-dependent TRIP6 condensation restrains Hippo hyperactivation and sustains cell fitness under osmotic stress

We next investigated the impact of TRIP6 phase separation on downstream Hippo pathway effectors under osmotic stress. Immunofluorescence staining showed that sorbitol treatment reduced the nuclear-to-cytoplasmic ratio of YAP in TRIP6-GFP cells (Figure S6A), while this effect was significantly exacerbated in TRIP6 ΔIDR2 transfected cells, indicating enhanced cytoplasmic retention of YAP (Figure 4A-B). Consistently, western blot revealed higher YAP Ser127 phosphorylation in TRIP6 ΔIDR2 cells than in WT cells under osmotic stress (Figures 4C, S6B). At the cellular level, clonogenic assays showed only a modest reduction in colony formation by TRIP6 ΔIDR2 cells relative to WT (Figure 4D-E). CCK-8 assays demonstrated that prolonged sorbitol treatment impaired viability more severely in TRIP6 ΔIDR2 cells than in WT cells (Figure 4F). To further confirm the difference in proliferation under osmotic stress, BrdU incorporation assays were performed. These assays showed that sorbitol treatment reduced BrdU incorporation and suppressed cell proliferation in TRIP6-GFP cells (Figure S6C), while this inhibitory effect was significantly exacerbated in TRIP6 ΔIDR2 cells, indicating further impaired proliferative capacity under osmotic stress (Figure 4G-H). Collectively, these data indicate that IDR2-mediated phase separation enables TRIP6 to buffer excessive Hippo pathway activation, thereby preserving cell proliferation and survival under osmotic stress. To validate our findings in a physiological tissue mechanics model, we employed K10-rtTA; TRE-Arhgef11^CA^ mice, in which elevated epidermal actomyosin contractility increases tissue tension and drives YAP1/TAZ-dependent basal stem cell hyperproliferation [27]. Immunofluorescence of E16.5 skin revealed prominent TRIP6 puncta in control epidermis but diffuse distribution upon Arhgef11 overexpression (Figure S6D). This *in vivo* observation is consistent with condensate disassembly seen in cells under osmotic stress, raising the possibility that TRIP6 condensates function as tension-responsive mechanosensitive assemblies.

**Figure 4.**
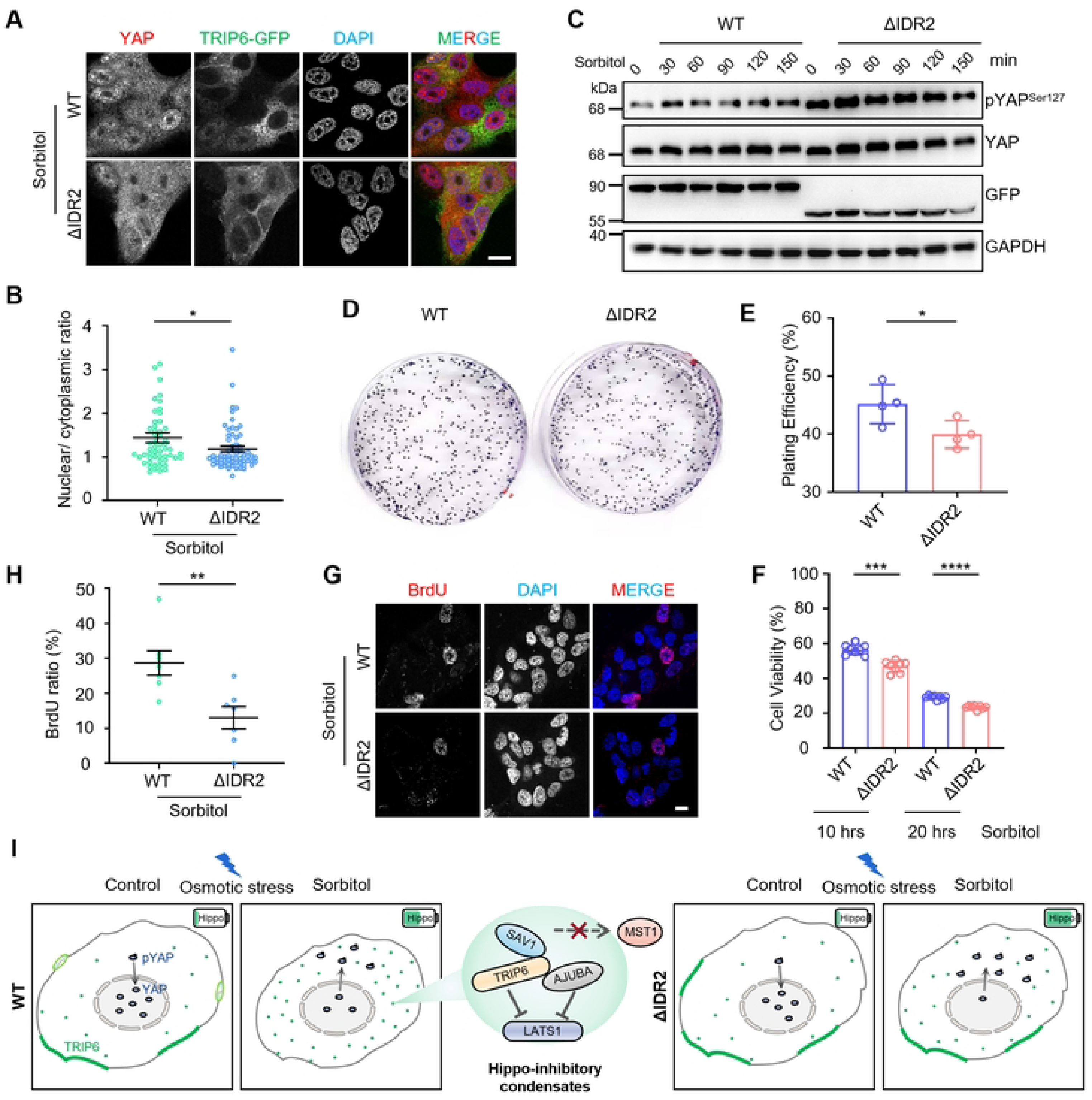
IDR2-dependent TRIP6 condensation restrains Hippo hyperactivation and sustains cell fitness under osmotic stress. (A and B) Immunofluorescence analysis of YAP subcellular localization in WT and ΔIDR2 cells under sorbitol-treated conditions. (A) Representative images. Scale bar, 10 μm. (B) Quantification of the nuclear-to-cytoplasmic YAP ratio showing that sorbitol-induced YAP cytoplasmic retention is significantly exacerbated in ΔIDR2 cells. n = 54 (WT) and 63 (ΔIDR2) cells from three independent experiments, respectively. Data are presented as mean ± SEM. Unpaired t test. *p < 0.05. (C) Western blot analysis of YAP phosphorylation at Ser127 in WT and ΔIDR2 cells treated with sorbitol for the indicated times, confirming higher YAP phosphorylation in ΔIDR2 cells under osmotic stress. (D and E) Clonogenic assays showing a modest reduction in colony formation by TRIP6-ΔIDR2 cells relative to WT. (D) Representative images of crystal violet-stained colonies. (E) Quantification of plating efficiency. n = 4 independent experiments per group. Data are presented as mean ± SD. Unpaired t test. *p < 0.05. (F) CCK-8 assay demonstrating that prolonged sorbitol treatment impairs viability more severely in TRIP6-ΔIDR2 cells than in WT cells. n = 8 independent experiments per group. Data are presented as mean ± SD. One-way ANOVA followed by Tukey’s multiple comparisons test. ***p < 0.001, ****p < 0.0001. (G and H) BrdU incorporation assays showing that TRIP6 ΔIDR2 cells exhibit significantly lower proliferative capacity than WT cells upon sorbitol treatment. (G) Representative images. BrdU (red), DAPI (blue). Scale bar, 10 μm. (H) Quantification of BrdU-positive cell ratio. n = 7 independent experiments per group, respectively. Data are presented as mean ± SEM. Unpaired t test. **p < 0.01. (I) Working model illustrating IDR2-mediated phase separation driving TRIP6 condensate redistribution under osmotic stress. The condensates selectively recruit the LATS1 inhibitor AJUBA and the MST1 scaffold SAV1 while excluding MST1 kinase, thereby exerting an inhibitory effect that constrains excessive Hippo pathway activation and preserves YAP nuclear localization.

Collectively, we demonstrate that TRIP6 is an intrinsically phase-separating LIM-domain scaffold whose IDR2-driven condensates redistribute rapidly under osmotic stress. By selectively recruiting AJUBA and SAV1 and strengthening LATS1 binding, these condensates act as a negative regulatory scaffold that constrains excessive Hippo activation to preserve cell proliferation and stress resilience (Figure 4I).

## Discussion

The Hippo pathway is a central regulator of cell proliferation and tissue homeostasis, and its dysregulation underlies tumorigenesis, organ fibrosis and defective tissue repair. In this study, we identify TRIP6 condensates as a negative regulatory node that buffers excessive Hippo pathway activation under hyperosmotic stress. This work complements the prevailing view of phase separation as a driver of Hippo activation by defining an inhibitory, mechanosensitive condensate that preserves signaling homeostasis under stress.

Membrane anchored cell-cell and cell-matrix adhesions can assemble phase separated condensates on the plasma membrane to support mechanotransduction. At focal adhesions, LIMD1 forms force dependent surface associated condensates, where TRIP6 functions as a recruited client protein [23]. At tight junctions, ZO-1 also builds mechanosensitive condensates at the plasma membrane to support junction maturation [28, 29]. Beyond structural roles, protein pools residing at cell junctions can dissociate and relocate to drive cytoplasmic signaling responses. This paradigm is exemplified by adherens junction resident β-catenin, which shuttles from cell contacts to the cytoplasm and nucleus to propagate Wnt signaling [30]. Drawing on these precedents, we hypothesise that TRIP6 may functionally switch from condensates restricted at adherens junctions to cytoplasmic signaling condensates upon hyperosmotic insult. The disassembly of TRIP6 condensates in a mouse model of elevated tissue tension further identifies them as mechanosensitive platforms linking tissue mechanics and Hippo-dependent growth control. This potential interconversion between junction tethered and cytosolic condensate states remains to be experimentally validated.

Multidomain LIM family proteins associated with cell junctions can employ intramolecular autoregulatory mechanisms. This regulatory mode is exemplified by LIMK1/2, which use a conserved PDZ domain surface for intramolecular autoinhibition independent of activation-loop phosphorylation [31]. Consistent with this, our data defines a bipartite regulatory logic governing TRIP6 condensate behaviour: the N-terminal IDR2 drives phase separation, while C-terminal LIM domains reduce condensate fluidity. TRIP6 condensates represent a distinct negative regulatory node within the Hippo phase separation network, contrasting with known activating condensates such as AMOT/KIBRA, MST1/2 and RIPK4-LATS assemblies [9, 11, 12]. This coexistence of activation and counterbalancing branches may reflect a general strategy for maintaining signaling homeostasis under stress. A second notable feature is the mechanosensitivity of TRIP6 condensates. Both hyperosmotic stress and elevated tissue tension *in vivo* trigger their disassembly, consistent with the role of TRIP6 in the α-catenin/vinculin tension-sensor complex [22] and suggesting a unified mechanotransduction axis linking mechanics, cytoskeletal remodeling, phase separation and Hippo output. Finally, the selective recruitment of AJUBA and SAV1 but not MST1 implies that TRIP6 condensates function as signal-sorting microdomains. By sequestering SAV1 away from MST1 while enriching the LATS inhibitor AJUBA [32, 33], these condensates may constrain Hippo activation at both upstream and downstream tiers. Future studies should address whether TRIP6 and activating condensates spatially coexist or compete, and whether AJUBA depletion abolishes the rheostat function of TRIP6 condensates.

Beyond osmotic stress alone, phase separation is increasingly recognized as a general mechanism for transducing osmotic signals across systems. In plants, molecular crowding drives reversible condensate assembly to transduce osmotic signals [34-36]. In mammalian cells, hyperosmotic stress elicits diverse adaptive responses through phase separation, including nuclear proteasome foci for ribosomal protein quality control [37] and BAG2-positive compartments for ubiquitin-independent protein degradation [38]. The osmolarity change further govern cell fate decisions, with acute osmotic shifts triggering apoptosis and gradual osmotic elevation favouring adaptive survival [39]. Within this stress-response landscape, Hippo signaling is no exception. Hyperosmotic stress triggers multiphase coalescence of functionally antagonizing Hippo-regulatory condensates to drive pathway activation, as exemplified by AMOT-KIBRA and SLMAP assemblies [12]. Notably, however, the Hippo-related examples documented so far predominantly illustrate how condensates initiate or amplify pathway activation. Our data reveal a complementary inhibitory signaling branch within this stress response network. Hyperosmotic stress triggers TRIP6 condensate redistribution to strengthen LATS1 binding. This change limits excessive Hippo activation and maintains cell proliferation and stress resilience. How TRIP6 condensates constrain Hippo signaling under physiological and pathological stress conditions warrants further investigation.

## Materials and Methods

### Cell culture and transfection

All cells including DLD1, HeLa, COS7 and HEK293T cells were cultured in Dulbecco’s modified Eagle’s medium (DMEM, Thermo Fisher) supplemented with 10% fetal bovine serum (ExCell) and 1% penicillin-streptomycin (Beyotime). All cells were cultured at 37 ° C in a humidified incubator with 5% CO_2_ .Cell transfections were performed using Lipo2000 transfection reagent (Thermo Fisher), HighGene plus transfection reagent (ABclonal), or PEI reagent (Polysciences) according to the manufacturer’s instructions. HEK293T, HeLa, and DLD1 cells were kindly provided by Prof. Jianwei Sun of Yunnan University. COS7 cells were kindly provided by Prof. Chonglin Yang of Yunnan University.

### Plasmids and constructs

Human TRIP6 cDNA was obtained from the previously constructed TRIP6-GFP plasmid, and all TRIP6 constructs below were cloned from this template [22]. For bacterial expression, TRIP6-WT and its truncation mutants, including N-terminal, C-terminal, ΔLIM3, ΔLIM2/3, ΔLIM1/2/3, ΔIDR1, ΔIDR2, and the FUS-IDR substitution mutant were cloned into pET-28a with an N-terminal His-GFP tag. Human AJUBA, SAV1 and MST1 cDNAs were likewise cloned into pET-28a with an N-terminal His-mCherry tag. For optoDroplet assays, TRIP6-WT and the same panel of truncation and FUS-IDR substitution mutants were cloned into the pCRY2-mCherry vector. For stable cell line generation, TRIP6-WT and TRIP6-ΔIDR2 were cloned into pLVX-Puro. For TurboID proximity labeling, TRIP6-WT and TRIP6-ΔIDR2 were cloned into a C-terminal TurboID-Flag (FT) vector, and free TurboID-FT was used as a negative control. For TRIP6 knockout, two TRIP6-targeting sgRNAs (5’-CCAGGGGGACCGGAGGATCG-3’ and 5’-ACGCTGGCCGAGCTGAATGG-3’) were cloned into the lentiCRISPR v2 plasmid. All constructs were verified by DNA sequencing.

### Generation of TRIP6 knockout and stable cell lines

All lentiviral particles were packaged in HEK293T cells by co-transfection of the corresponding transfer plasmid with packaging plasmids psPAX2 and pMD2G using PEI reagent. Viral supernatants were harvested 48-72 h post-transfection, filtered, and applied to target cells supplemented with 10 μg/mL polybrene (Biosharp) to enhance transduction efficiency. TRIP6-knockout DLD1 cells were generated by infection with lentivirus carrying the lentiCRISPR v2 construct encoding the two TRIP6-targeting sgRNAs described above. Positive cells were selected with puromycin (Solarbio) and maintained under continuous antibiotic pressure. For TRIP6 reconstitution, TRIP6-WT-GFP and TRIP6-ΔIDR2-GFP were cloned into the pLVX lentiviral vector, packaged into lentivirus, and transduced into TRIP6-knockout DLD1 cells. GFP-positive cells were enriched by flow sorting on a BD FACSAria SORP cell sorter. For TurboID proximity labeling assays, TRIP6-WT-FT and TRIP6-ΔIDR2-FT lentiviruses were generated from the C-terminal TurboID-Flag (FT) transfer vector and transduced into DLD1 cells. Positive cells were selected with puromycin and maintained under continuous antibiotic pressure.

### Antibodies

Primary antibodies used in this study included rabbit anti-TRIP6 (Proteintech, 21163-1-AP, 1:2,000 for WB, 1:200 for IF), rabbit anti-LATS1 (Proteintech, 17049-1-AP, 1:200 for IF), rabbit anti-YAP (HUABIO, ET1608-30, 1:1,000 for WB, 1:100 for IF), rabbit anti-phospho-YAP (Ser127) (CST, 4911, 1:1,000 for WB), mouse anti-Flag (Sigma, F1804, 1:4,000 for WB), mouse anti-BrdU (Proteintech, 66241-1-Ig, 1:300 for IF), mouse anti-GFP (Proteintech, 66002-1-Ig, 1:20,000 for WB), mouse anti-GAPDH (Proteintech, 60004-1-Ig, 1:50,000 for WB), DAPI (MeilunBio, MA0127, 1:500), 594-Phalloidin (Proteintech, PF00003, 1:400), 488-Phalloidin (Proteintech, PF00001, 1:400), 594-Streptavidin (Yeasen, 35107ES60, 1:500) and HRP-conjugated streptavidin (Beyotime, A0303, 1:4,000 for WB). Secondary antibodies included HRP-conjugated goat anti-rabbit and anti-mouse IgG (CST, 1:4,000 for WB), and CoraLite 488/594-conjugated goat anti-rabbit, anti-mouse IgG (Proteintech, 1:500 for IF).

### Co-immunoprecipitation and western blot

For co-immunoprecipitation assays, HEK293T cells were cultured in 10-cm dishes. Cells were harvested and resuspended in ice-cold lysis buffer containing 50 mM Tris-HCl (pH 7.4), 150 mM NaCl, 1 mM EDTA, 1 mM DTT, 0.1% Triton X-100 and 5% glycerol, supplemented with 1 mM PMSF (Solarbio). Cell lysates were prepared by sonication, followed by centrifugation at 12,000 rpm for 20 min at 4 ° C. The supernatants were collected and incubated with anti-Flag agarose beads (Yeasen, 20585ES08) for 4 h at 4 °C. After incubation, the beads were washed three times with lysis buffer, boiled in 1 × SDS sample buffer for 10 min at 100 ° C, and then subjected to western blot analysis. For western blot, proteins were separated by SDS-PAGE and transferred to PVDF membranes. Membranes were blocked with 5% non-fat milk in PBST for 1 h at room temperature and incubated with primary antibodies overnight at 4 ° C, followed by HRP-conjugated secondary antibodies for 1 h. Signals were detected using an enhanced chemiluminescence system.

### Protein expression and purification

Recombinant proteins were expressed in *Escherichia coli* BL21 cells. Bacteria were grown at 37℃ to OD_600_=0.8 and induced with 1 mM IPTG (OriLeaf) at 16°C for 20 h. Cells were harvested by centrifugation and lysed by sonication in lysis buffer (50 mM Tris-HCl, pH 8.0, 300 mM NaCl, 10 mM imidazole, 1 mM PMSF). Lysates were clarified by centrifugation at 8,000 rpm for 30 min, and proteins were purified using Ni-NTA agarose beads (Yeasen, 20502ES60). Proteins were eluted with elution buffer (50 mM Tris-HCl, pH 8.0, 150 mM NaCl, 250 mM imidazole). Protein purity was assessed by SDS-PAGE and Coomassie Blue staining.

### *In vitro* phase separation assay

Purified proteins were diluted from stock solutions into phase separation buffer (25 mM Tris-HCl, pH 8.0, 150 mM NaCl) to reach the indicated final concentrations. For concentration-dependent assays, TRIP6 was serially diluted to final concentrations of 10, 20, 30 and 40 μM. For ionic strength-dependent assays, TRIP6 was set at a final concentration of 30 μM, and NaCl concentrations were adjusted to 50, 150, 300 or 600 mM using a 2 M NaCl stock solution.

Protein solutions were transferred to glass-bottom dishes and imaged on a Zeiss LSM800 or LSM980 confocal microscope equipped with a Plan-Apochromat 63×/1.4 oil immersion objective. Droplet area was quantified using Fiji/ImageJ software. For *in vitro* co-phase separation assays, 20 μM purified TRIP6-GFP was mixed with 10 μM His-mCherry-AJUBA, His-mCherry-SAV1, or His-mCherry-MST1 in phase separation buffer. Samples were imaged by confocal microscopy, and fluorescence intensity line scans across individual droplets were generated using Fiji/ImageJ software.

### TurboID proximity labeling and mass spectrometry

DLD1 cells stably expressing TRIP6-WT-FT, TRIP6-ΔIDR2-FT, or free TurboID-FT were cultured to 80% confluence in 10-cm dishes. For biotin labeling, cells were incubated with biotin (Sigma, B4501) in complete medium for 2 h at 37 ° C. Cells were resuspended, lysed and centrifuged to collect supernatants as described above. Free biotin was removed by ultrafiltration column (Pall, Microsep MCP003C, 4-5 mL). Biotinylated proteins were enriched by incubation with streptavidin agarose resin (Yeasen, 20512ES08) at room temperature for 1 h on a rocking platform followed by three times washing with PBST. Bound proteins were eluted by boiling in 1 × SDS sample buffer at 95°C for 10 min. Eluted proteins were separated by 10% SDS-PAGE, and protein bands were excised for in-gel trypsin digestion. MS analysis was performed at the Mass Spectrometry Facility of Yunnan University. Protein identification and label-free quantification were performed using MaxQuant software (version 1.6.4.0). Candidate proteins were selected based on log_2_ intensity ratio and number of unique peptides enriched in TRIP6-WT samples relative to control and TRIP6-ΔIDR2 groups. Hierarchical clustering heatmap, principal component analysis and KEGG pathway enrichment analysis were performed using the Servicebio online multi-omics analysis platform (https://multi-omics.servicebio.cn/).

### CCK-8 assay

Cell viability was assessed using the Cell Counting Kit-8 (CCK-8) assay (Glpbio, GK10001). Briefly, cells were seeded into 96-well plates at 10^4^-10^5^ cells per well in 100 μL complete medium and cultured overnight. After treatment with or without 0.4 M sorbitol for the indicated time periods, 10 μL of CCK-8 solution was added to each well, followed by incubation for 2 h at 37℃. The optical density (OD) was measured at 450 nm using a microplate reader. Cell viability was calculated relative to the control group.

### Colony formation assay

Cells were seeded in 3-cm dishes at 500 cells per dish and cultured for 12 days. Colonies were fixed with 4% paraformaldehyde for 10 min and stained with a solution comprising 0.5% crystal violet (OriLeaf). Colony images were captured, and colony area was quantified using Fiji/ImageJ software.

### BrdU incorporation assay

Cells were treated with 0.4 M sorbitol and 30 μg/mL BrdU simultaneously for 1.5 h before fixation. After 4% paraformaldehyde fixation and 0.2% Triton X-100 permeabilization, DNA was denatured with 2 N HCl at 37°C for 30 min, followed by two washes with PBS. Cells were then incubated with anti-BrdU antibody, followed by 594-conjugated secondary antibody and DAPI. The percentage of BrdU-positive cells was quantified from at least 100 cells per condition across three independent experiments.

### Mouse experiments and Tissue processing

K10-rtTA; TRE-Arhgef11^CA^ mice [27] were maintained in a barrier facility under a 12-h light/dark cycle with free access to food and water. To induce Arhgef11 expression in differentiated epidermal cells, pregnant females were fed doxycycline water from embryonic day 13.5 (E13.5) until harvest. Embryos were collected at E16.5 and embedded in OCT compound. Frozen sections were cut using a cryostat for subsequent experiments. All animal experiments were approved by the Institutional Animal Care and Use Committee of Yunnan University.

### Immunofluorescence staining

For cultured cells grown on glass coverslips, samples were fixed with 4% paraformaldehyde for 10 min at room temperature, permeabilized with 0.1% Triton X-100 in PBS for 2 min, and blocked with 1% BSA in PBS for 30 min. Cells were then incubated with primary antibodies for 2 h at room temperature or overnight at 4°C, followed by 488/594-conjugated secondary antibodies and DAPI for 1 h. For F-actin staining, 594-phalloidin was included during the secondary antibody incubation. Coverslips were mounted with anti-fade mounting medium (UElandy). For tissue sections, permeabilization was performed with 0.2% Triton X-100 in PBS for 10 min, and blocking was extended to 1 h at room temperature with 3% BSA in PBS. Sections were incubated with anti-TRIP6 antibody overnight at 4°C, followed by 488-conjugated secondary antibody, 594-phalloidin and DAPI for 1 h at room temperature. Slides were mounted with anti-fade mounting medium (Solarbio).

### FRAP analysis

FRAP experiments were performed using Zeiss LSM 980. Selected regions within TRIP6 condensates were photobleached with 100% laser power using a 488-nm laser. Time-lapse images were acquired at 5-s intervals. Fluorescence intensities of regions of interest were normalized to the initial intensity before bleaching. FRAP data were analyzed using GraphPad Prism.

### OptoDroplet assay

COS-7 cells expressing CRY2-mCherry-TRIP6 constructs were seeded on glass-bottom dishes. At 24 h post transfection, live cells were imaged using LSM 800 with a 63× oil objective. Cells expressing optoDroplet plasmids at comparable levels were selected for light-induced activation with a 488-nm laser. mCherry was excited and visualized using a 594-nm laser. Time-lapse images were acquired every 5 s for at least 150 s to monitor condensate formation.

### Structured illumination microscopy imaging

Structured illumination microscopy was performed using a Zeiss Elyra 7 microscope equipped with Lattice SIM^2^ module (Zeiss, Jena, Germany) and a 63×/1.4 NA oil objective. Serial Z-stacks were acquired at 0.1 μm axial intervals. Following image acquisition, SIM super-resolution reconstruction was performed. Z-stack images were subsequently generated in Fiji/ImageJ, from which maximum-intensity projections were produced.

### Quantification and statistical analysis

All quantitative data are presented as mean ± standard deviation (SD) or standard error of the mean (SEM) from at least three independent experiments. Statistical analyses were performed using GraphPad Prism 8 software. Comparisons between two groups were made using unpaired two-tailed Student’s t-test. Comparisons among multiple groups were made using one-way analysis of variance (ANOVA) followed by Tukey’s multiple comparisons test or Dunnett’s multiple comparisons test selected according to the experimental design (as specified in individual figure legends). A P-value less than 0.05 was considered statistically significant.

## Author Contribution

Conceptualization: Weihong Fu, Wenxiu Ning.

Formal analysis: Weihong Fu, Wenxiu Ning.

Funding acquisition: Wenxiu Ning.

Investigation: Weihong Fu, Yuxi Yang.

Methodology: Weihong Fu, Hua Li.

Resources: Wenxiu Ning.

Supervision: Wenxiu Ning.

Validation: Weihong Fu, Yuxi Yang, Lin Xie, Junying Li.

Visualization: Weihong Fu.

Writing - original draft: Weihong Fu, Hua Li, Wenxiu Ning.

Writing - review & editing: Weihong Fu, Lin Xie, Hua Li, Wenxiu Ning.

All authors read and approved the final version of the manuscript.

## Data availability statement

All relevant data are within the paper and its Supporting Information files. The raw mass spectrometry (MS) data have been deposited to the ProteomeXchange Consortium (https://proteomecentral.proteomexchange.org) via the iProX partner repository [40, 41] with the dataset identifier PXD085596. All data generated in this study are available from the corresponding author upon reasonable request.

## Acknowledgments

We are grateful to Professor Xuna Wu for mass spectrometry analysis, the core facilities of Yunnan University for imaging. All animal studies were approved by the Yunnan University Institutional Animal Care and Use Committee (IACUC, approval no. YNU20240797), and performed in accordance with institutional and national guidelines.

## Supporting information

**Figure S1. TRIP6-GFP undergoes condensate redistribution under osmotic stress.**

(A) Western blot validation of TRIP6-GFP expression in the TRIP6 KO background. (B) Immunofluorescence staining of TRIP6-GFP cells under basal conditions, 0.4 M sorbitol and 0.2 M NaCl treatment, showing that TRIP6-GFP faithfully recapitulates the osmotic stress-induced redistribution of endogenous TRIP6. Scale bar, 10 μm (white) and 2 μm (yellow). (C) Coomassie blue staining of purified recombinant His-GFP-TRIP6 protein analyzed by SDS-PAGE.

**Figure S2. TRIP6 condensate formation is independent of its LIM domains.**

(A) Coomassie blue staining of purified full-length TRIP6 (WT), N-terminal (N) and C-terminal (C) fragments. (B) Coomassie blue staining of purified WT and sequential LIM deletion mutants (ΔLIM3, ΔLIM2/3, ΔLIM1/2/3). (C and D) *In vitro* phase separation of WT and LIM deletion mutants at 40 μM in 150 mM NaCl. (C) Representative images. Scale bar, 5 μm. (D) Quantification of droplet area showing no significant difference among genotypes. n = 80 droplets for each group. Data are presented as mean ± SEM. One-way ANOVA followed by Tukey’s multiple comparisons test. ns, not significant, **p < 0.01.

**Figure S3. ΔIDR2-FUS chimera rescues TRIP6 condensation.**

(A) Coomassie blue staining of purified WT, ΔIDR1 and ΔIDR2 proteins. (B) Domain architecture of TRIP6 WT, ΔIDR2 and the ΔIDR2-FUS chimera in which IDR2 is replaced by the FUS IDR. (C) OptoDroplet assay of WT, ΔIDR2 and ΔIDR2-FUS showing that FUS-IDR substitution restores light-induced condensate formation. Scale bar, 10 μm. (D) Coomassie blue staining of purified WT, ΔIDR2 and ΔIDR2-FUS proteins. (E and F) *In vitro* phase separation of WT, ΔIDR2 and ΔIDR2-FUS at 10 μM in 50 mM NaCl. (E) Representative images showing that the ΔIDR2-FUS chimera forms large droplets at low protein concentration. Scale bar, 10 μm. (F) Quantification of droplet area. n = 185 (WT), 161 (ΔIDR2), 89 (ΔIDR2-FUS) droplets, respectively. Data are presented as mean ± SEM. One-way ANOVA followed by Tukey’s multiple comparisons test. ****p < 0.0001.

**Figure S4. Validation of TurboID labeling.**

(A) Streptavidin pull-down and western blot analysis confirming expression of free TurboID-FT (FT), TRIP6-WT-FT and TRIP6-ΔIDR2-FT, and efficient biotinylated protein enrichment across all three groups. (B) Immunofluorescence staining showing that TRIP6-WT-FT directs biotin deposition to discrete cytoplasmic puncta matching TRIP6 condensate localization, whereas TRIP6-ΔIDR2-FT produces diffuse cytoplasmic biotin signals. Scale bar, 10 μm. (C) Principal component analysis of the quantitative proteomic dataset showing clear separation among control, TRIP6-WT and TRIP6-ΔIDR2 samples with excellent intra-group reproducibility. (D) Coomassie blue staining of purified His-mCherry-tagged AJUBA, SAV1 and MST1 proteins.

**Figure S5. IDR2 enhances the TRIP6-LATS1 colocalization under osmotic stress.**

(A and B) Flow cytometry analysis and gating for enrichment of (A) TRIP6-WT-GFP and (B) TRIP6-ΔIDR2-GFP positive cells. (C-G) Immunofluorescence analysis of TRIP6-LATS1 co-localization in WT and ΔIDR2 cells under basal and sorbitol-treated conditions. (C) Representative images showing that sorbitol enhances co-localization between TRIP6-WT and LATS1 but not between TRIP6-ΔIDR2 and LATS1. TRIP6-GFP (green), LATS1 (red), DAPI (blue). Scale bar, 10 μm (white) and 2 μm (yellow). (D-G) Fluorescence intensity line-scan profiles across individual cells for WT under control (D) and sorbitol (E) conditions, and for ΔIDR2 under control (F) and sorbitol (G) conditions. F.I., fluorescence intensity.

**Figure S6. TRIP6 condensates constrain Hippo hyperactivation.**

(A) Sorbitol treatment induces YAP cytoplasmic retention with reduced nuclear localization. Representative immunofluorescence images showing YAP (red), TRIP6-GFP (green) and DAPI (blue) staining under control and sorbitol-treated conditions. Scale bar, 10 μm. (B) Western blot analysis confirming higher YAP Ser127 phosphorylation in TRIP6 ΔIDR2 cells than in WT cells under sorbitol treatment. (C) Sorbitol treatment reduces BrdU incorporation and suppresses cell proliferation. Representative immunofluorescence images showing BrdU (red) and DAPI (blue) staining in TRIP6-GFP cells under control and sorbitol-treated conditions. Scale bar, 10 μm. (D) Immunofluorescence staining of E16.5 embryonic skin sections from control and K10-rtTA; TRE-Arhgef11^CA^ mice. TRIP6 (green) forms prominent puncta in control epidermis but becomes markedly diffuse in the Arhgef11-overexpressing epidermis with elevated tissue tension. F-actin (red), DAPI (blue). Scale bars, 20 μm (white) and 2 μm (yellow).

**S1 Raw images.** PDF file containing un-cropped images of all western blots and gels in this manuscript.

